# The representational geometry for naturalistic textures in macaque V1 and V2

**DOI:** 10.1101/2024.10.18.619102

**Authors:** Abhimanyu Pavuluri, Adam Kohn

## Abstract

Our understanding of visual cortical processing has relied primarily on studying the selectivity of individual neurons in different areas. A complementary approach is to study how the representational geometry of neuronal populations differs across areas. Though the geometry is derived from individual neuronal selectivity, it can reveal encoding strategies difficult to infer from single neuron responses. In addition, recent theoretical work has begun to relate distinct functional objectives to different representational geometries. To understand how the representational geometry changes across stages of processing, we measured neuronal population responses in primary visual cortex (V1) and area V2 of macaque monkeys to an ensemble of synthetic, naturalistic textures. Responses were lower dimensional in V2 than V1, and there was a better alignment of V2 population responses to different textures. The representational geometry in V2 afforded better discriminability between out-of-sample textures. We performed complementary analyses of standard convolutional network models, which did not replicate the representational geometry of cortex. We conclude that there is a shift in the representational geometry between V1 and V2, with the V2 representation exhibiting features of a low-dimensional, systematic encoding of different textures and of different instantiations of each texture. Our results suggest that comparisons of representational geometry can reveal important transformations that occur across successive stages of visual processing.

---

Historically, the study of how visual information is represented at different stages of cortical processing—and inferences about the computations that create those representations—has been based on measuring single neuron selectivity for different images. This approach has provided much of our knowledge of visual cortical function (Hubel and Wiesel, 1962; Movshon et al., 1978; Desimone et al., 1984). However, it has struggled to elucidate representations in many mid-level and higher visual areas (Boynton and Hegde, 2004, Pasupathy et al., 2020).

An alternative approach to understanding visual representations is to study how they manifest in neuronal populations, more specifically in the geometry of their responses (Chung and Abbott, 2021; Langdon et al., 2023). Visual representational geometry refers to the distribution of activity in the neuronal population response space generated by different images, also referred to (loosely) as a response ‘manifold’. Though a complete understanding of the selectivity of each single neuron would allow one to derive the representational geometry, in practice our understanding of single neuron selectivity is insufficient to predict the geometry, even in V1 (Stringer et al., 2019, Kriegeskorte and Wei, 2021). Instead, characterizing the representational geometry requires analyzing the responses of many cells together to an ensemble of many images.

What is known about the representational geometry in the visual cortex? Prior work has described the dimensionality of population representation in areas such as V1 (Cowley et al., 2016; Stringer et al., 2019) and IT (Lehky et al., 2014) and across visual areas of the mouse (Frourdarakis et al., 2021). Dimensionality is a key measure of the representational geometry, but of course this measure alone paints an incomplete picture. In addition, since each study has used a distinct stimulus ensemble and focused on a single brain area, it has remained unclear how representations evolve across stages of processing (but see Frourdarakis et al., 2021). Comparisons across areas provide insight into which aspects of geometry are emphasized as processing progresses.

Theoretical and computational work has begun to explore the functional advantages and disadvantages of different representational geometries. In general, lower-dimensional representations are more robust to noise and can afford better generalization performance (Stringer et al., 2019; Bernardi et al., 2020; Johnston and Fusi, 2023), whereas higher dimensional representations are often advantageous for encoding capacity (Babadi and Sompolinsky, 2014; Stringer et al., 2019; Cohen et al., 2020) and can support few-shot learning (Sorscher et al., 2022). However, these functional measures—encoding capacity, robustness, generalization, learning, and others—are not fully specified by the response dimensionality. They depend also on additional features of the geometry.

Here, we analyze neuronal population responses recorded in areas V1 and V2 of macaque monkeys to a range of naturalistic textures. We focus on naturalistic textures because of previously reported differences in V1 and V2 single neuron responses to these stimuli (Freeman et al., 2013; Ziemba et al., 2016, 2019). We assess how the representational geometry for these images differs between V1 and V2. We find that V2 population responses to textures are lower dimensional than in V1; control experiments with gratings, in contrast, revealed V2 responses are higher dimensional than in V1. To understand better the nature of the representational geometry of textures, we measured the alignment of different components of the response manifolds and found that responses to different instantiations of a texture were aligned to the manifold representing different textures. Together, these features suggest the construction of a systematic, low-dimensional embedding of texture stimuli. We show the representational geometry in V2 allows for better discrimination between out-of-set, novel textures. Finally, we find a divergence between the representational geometry in cortex and those of common convolutional networks, indicating that measures of population geometry can reveal different encoding schemes in brain and machine.

## Results

We recorded neurons in V1 and V2 simultaneously, using up to four Neuropixel probes inserted normal to the cortical surface (Figure 1A). Using the well-described retinotopy of V1 and V2, we targeted our electrode penetrations to provide neurons in the two areas with overlapping or nearby spatial receptive fields. We first measured the spatial receptive fields of the sampled neurons using a sparse mapping protocol. Figure 1B shows the receptive field maps for an example probe, with V1 and V2 receptive fields evident in different sections of the probe. We then centered our stimuli on the sampled population, using a stimulus size sufficient to cover all receptive fields. Receptive fields were 1-5 degrees eccentric, in the lower visual field. We analyzed the response of 167 V1 neurons and 272 V2 neurons that met inclusion criteria, from four anesthetized monkeys (see Methods).

**Figure 1:**
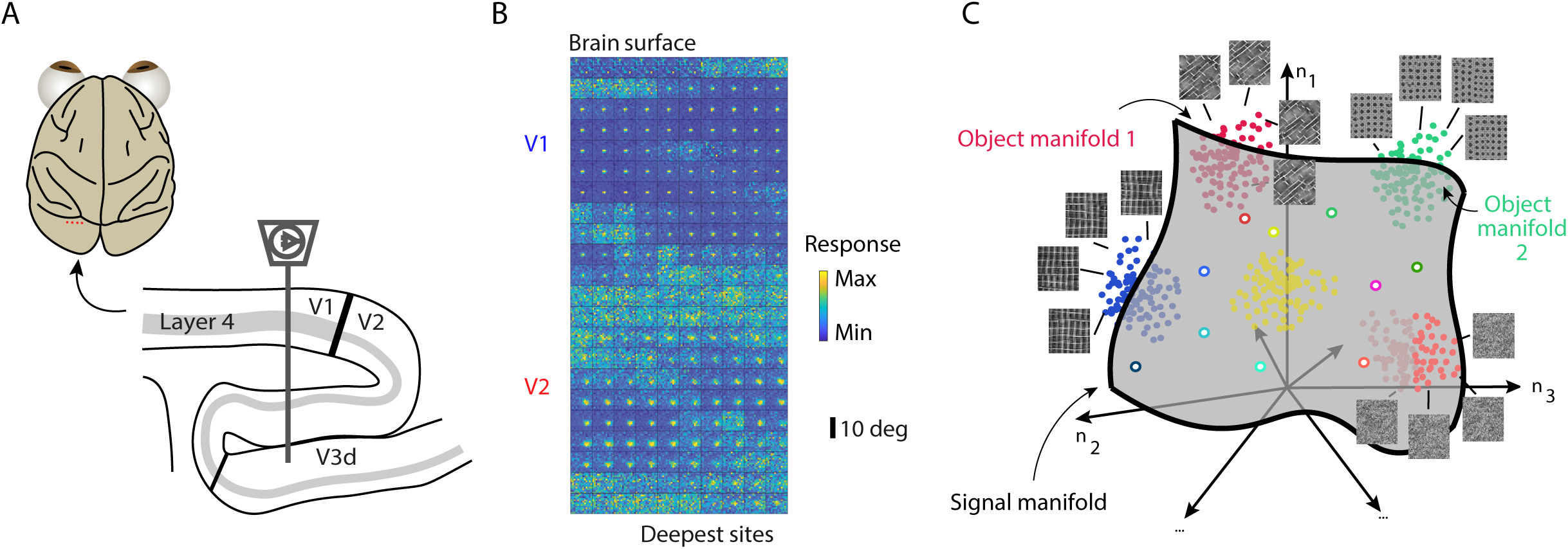
Experimental design. (**A**) Recording approach. Dorsal view of the macaque brain; electrode insertion sites indicated with orange dots just behind the lunate sulcus. Sagittal section of occipital cortex showing how V1 and V2 were sampled with a single probe. (**B**) Example spatial receptive field mapping data from one probe in one recording session. Each subplot shows the receptive field measured at one recording site, with the color indicating the activity measured for small gratings presented at a particular position. Sites are arranged based on their position on the probe, from the most dorsal at the top left to the most ventral at the bottom right. (**C**) The population representational geometry. The signal manifold is defined by the population responses to different texture families (indicated open circles of different colors, indicating the mean of the respective set of points); the object manifold is defined by the responses to multiple samples of a particular texture (indicated by filled circles of a particular color).

We measured responses in V1 and V2 to naturalistic textures, synthesized with the Portilla and Simoncelli algorithm (Portilla and Simoncelli, 2000). This algorithm summarizes a source image with a set of statistics based largely on responses of linear filters of different sizes, orientations, and spatial frequencies applied to that image, as well as the relationships of the filter responses to each other. To synthesize textures, patches of random pixels are iteratively transformed until their statistics match those of the original image, though the synthesized and original images will differ at the pixel level. We presented both synthetic textures derived from different source images (termed texture families) and different instantiations of each of those textures (termed samples; Freeman et al., 2013). Each sample within a texture family had the same statistics but was synthesized from a different noise seed.

We presented texture families and samples in two stimulus ensembles. One ensemble consisted of 90 texture families with 10 samples of each family; each distinct image was presented 10 times. This ensemble was used to measure the ‘signal manifold’ in the population response space, defined using the mean responses of each neuron across samples and presentations (Figure 1C, open circles). The second ensemble consisted of 10 texture families, with 90 samples of each, also each presented 10 times. We defined the trial-averaged population responses to the samples of each texture as an ‘object manifold’ (Figure 1C; filled circles; 5 different object manifolds indicated with the different colors). The texture families used in the signal manifold and object manifold ensembles were distinct from each other.

We presented the same ensembles in each animal. Since our analysis focused on trial-average responses, we combined all recorded units in each area into a ‘pseudo-population’ and performed our analyses on different random selections of units within these sets.

### Response dimensionality

We first characterized the dimensionality of the signal manifold, in V1 and V2. We performed principal component analysis on the measured population responses, after Z-scoring the response of each neuron across textures. Z-scoring ensured that the structure identified would not be unduly swayed by the responses of any high-variance units.

Figure 2A shows example eigenspectra for several randomly selected subsets of 50 neurons in V1 (blue) and V2 (red). The distribution of eigenvalues in V2 tended to have a stronger contribution from the first few eigenvectors (principal components) than the corresponding data in V1. To quantify these spectra, we calculated the participation ratio: a common summary statistic of the eigenspectrum, and a standard measure of dimensionality (Mazzucato et al., 2016; Litwin-Kumar et al., 2017). If the spectrum is flat, as for a distribution in which each neuron’s response is independent and of equal variance, then the participation ratio will equal the number of neurons. If the spectrum is dominated by a single component, then the participation ratio will approach a value of 1.

**Figure 2:**
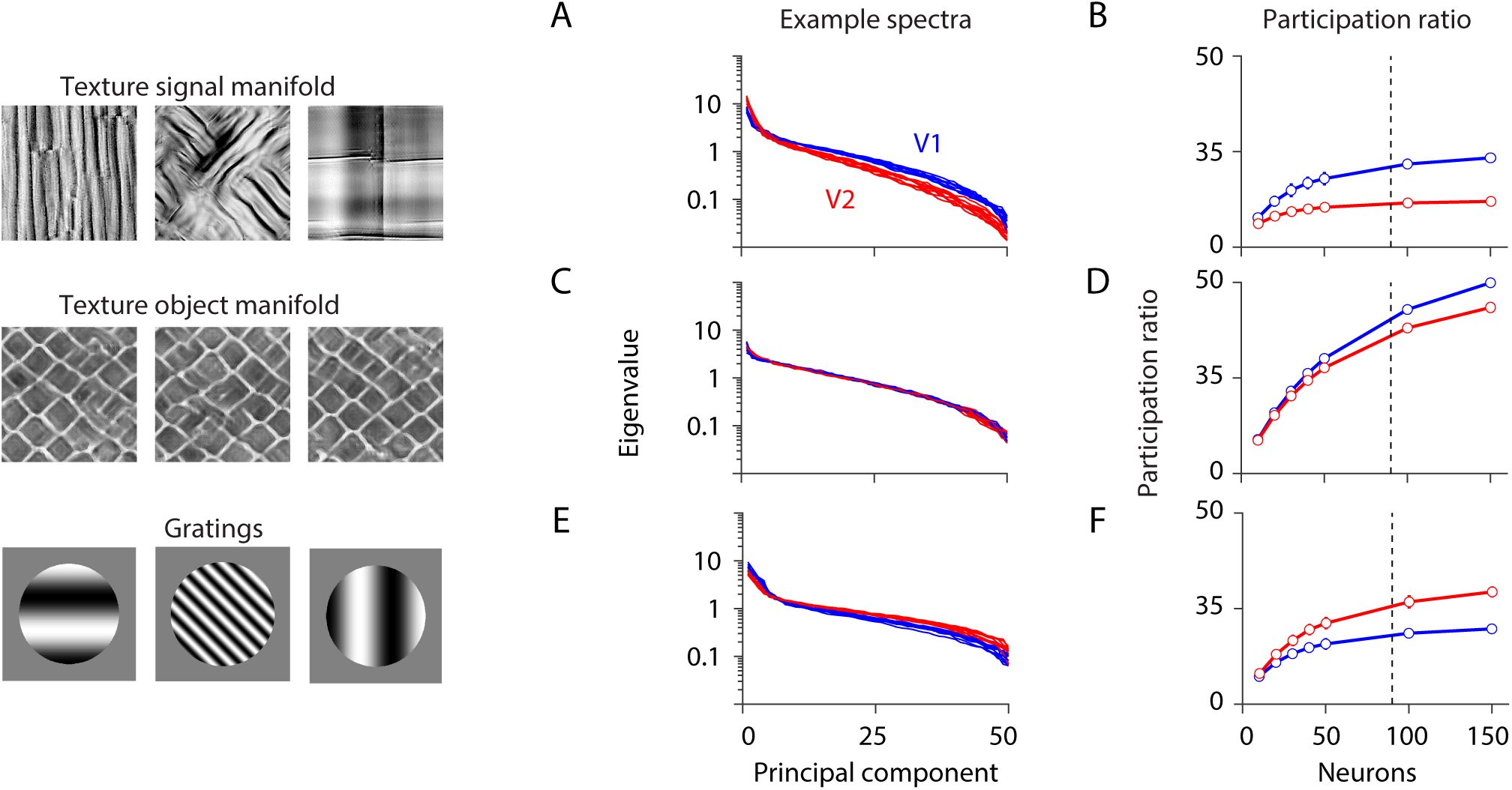
Participation ratio for different stimulus ensembles. (A) Example eigenspectra for V1 (blue) and V2 (red) population responses to the texture signal manifold ensemble. (B) The participation ratio in V1 and V2, for neuronal populations of different sizes. Dashed vertical line indicates the population size at which the number of sampled neurons matches the number of stimuli in the ensemble (n=90). (C) Example spectra for responses to different samples of a texture object manifold stimulus ensemble. (D) The participation ratio for population responses to texture object manifold ensemble. (E) Example spectra for responses to ensembles of sinusoidal gratings of different orientations and spatial frequencies. (F) Participation ratio for responses to gratings.

We calculated the participation ratio for different population sizes, ranging from 5 to 150 neurons, in which each population was created by sampling from the pool of neurons without replacement. We note that the signal manifold was measured using responses to 90 texture families—thus, for the larger population sizes, the number of observations was smaller than the number of recorded neurons, capping the observable dimensionality.

Consistent with the example eigenspectra, the participation ratio was lower in V2 (Figure 2B, red) than V1 (blue) for each population size. For populations of 50 neurons, for instance, the participation ratio was 10.6±1.2 (SD) in V2 and 18.2±1.9 in V1 (p<0.001 for difference between areas; bootstrap test, see Methods). The differences were more apparent for larger populations but were statistically distinguishable from chance for each population size, even those as small as 10 neurons (p<0.001; bootstrap test). Thus, V2 population responses to ensembles of different textures are lower dimensional than the responses in V1.

We performed similar analyses on responses to our second stimulus ensemble—the object manifolds—consisting of many instantiations (samples) of a few texture families. We computed the participation ratio for responses to each texture family separately. Participation ratios were higher for these responses (Figure 2D) than for those to the signal manifold ensemble, indicating higher-dimensional response distributions. This is expected: different texture families should yield more structured population responses than different samples of the same family. Nevertheless, response dimensionality was again lower dimensional in V2 than V1 (Figure 2C,D; for 50 neurons, 27.7±0.5 vs. 30.2±0.3, SEM across 10 object manifolds, p=0.02 for difference between the two, Wilcoxon rank sum test), particularly for larger populations.

To test whether population responses in V2 are consistently lower dimensional than those in V1, we presented an additional ensemble of images—consisting of gratings of many different orientations and spatial frequencies (see Methods)—in a subset of sessions (191 cells in V1; 235 in V2). Notably, population responses to gratings were higher dimensional in V2 than V1 (Figure 2E,F; for 50 neurons, 21.48±1.6 SD vs. 15.6±1.5; p<0.001, bootstrap test). Note that our comparison is not across images classes within a population—for example, V1 dimensionality for gratings versus textures, for which differences could trivially reflect the differing complexity of different stimulus ensembles—but rather a comparison across populations for a given image ensemble (V1 vs V2 for either gratings or textures).

We wondered whether the dimensionality differences in population responses to gratings and textures in V1 and V2—V2 responses are lower dimensional than V1 for textures but higher dimensional for gratings—might be due to a subset of images. To test this possibility, we randomly selected many different subsets of 60 images from each ensemble and calculated the participation ratio for 40 neurons. We chose these numbers to strike a balance between having enough images and cells to capture the relevant structure, while attempting to reduce the overlap between the different random subsamples. Figure 3 shows the participation ratio in V1 and V2 for responses to the two types of images. In nearly every case, V2 dimensionality is lower than that in V1 for responses driven by textures (black; 9.7±1.4 (S.D.) vs. 15.5±1.6, p<0.001, paired t-test); in contrast, V2 dimensionality is higher than V1 for responses driven by gratings (gray; 17.6±1.5 vs. 13.6±1.3; p<0.001, paired t-test).

**Figure 3:**
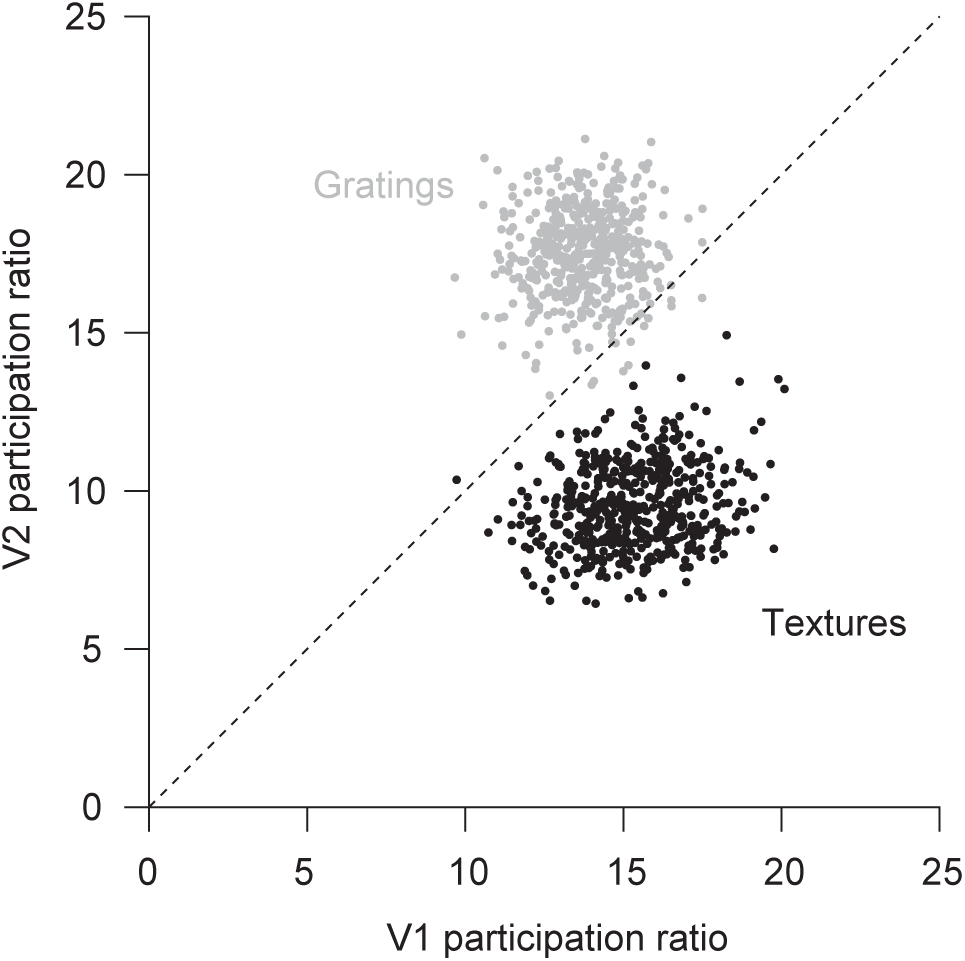
Comparison of the participation ratio for V1 (abscissa) and V2 (ordinate) population responses, for gratings (gray) and signal manifold textures (black). Each circle corresponds to a different random subsample of 40 neurons responding to a different subset of 60 images in the full ensemble.

In complementary analysis, we characterized the representational geometry in V1 and V2 using factor analysis (Yu et al., 2009; Williamson et al., 2016). Factor analysis is similar in spirit to PCA but relies on a probabilistic model that partitions the measured variance into a component that is shared across neurons and another which is private to each neuron. Consistent with the view provided by PCA, we found greater shared variance in V2 responses to textures than in V1, and less shared variance in V2 responses to gratings than in V1 (Supplementary Figure 1).

We conclude that the population representation of ensembles of different texture families and different texture samples is lower dimensional in V2 than V1. The reduction in representational dimensionality is not evident in population responses to gratings. This dependence on stimulus type indicates that differences in dimensionality do not simply reflect fixed, gross differences between these areas such as the number of neurons available to encode visual stimuli or consistent, strong differences in responsivity or sparsity.

### Alignment of manifolds

Low dimensional representations can facilitate generalization and robustness to noise (Stringer et al., 2019; Bernardi et al., 2020; Johnston and Fusi, 2023). But the degree to which this is the case will depend on how the low dimensional representation is constructed, namely on whether there is a systematic and orderly mapping of inputs onto the primary axes of the space. To understand better the representational geometry for textures in V1 and V2, we measured the alignment of the object manifolds to each other and to the signal manifold. We reasoned that if there was a systematic embedding of different texture images in a low-dimensional response space, than different manifolds should have some relationship to each other.

To quantify alignment, we calculated an index which measures how much response variance for one condition (e.g. data B) is captured by a set of dimensions identified using responses to a different condition (data A), relative to the variance captured by the optimal dimensions (i.e., the principal components of responses to data B; Elsayed et al., 2016). If two distributions have similar structure, the dimensions identified from one distribution will capture much of the structure of the other distribution, resulting in an alignment score near 1 (Figure 4A, top). If the two distributions have different structures, little variance will be captured, and the alignment score will approach zero (Figure 4A, bottom).

**Figure 4:**
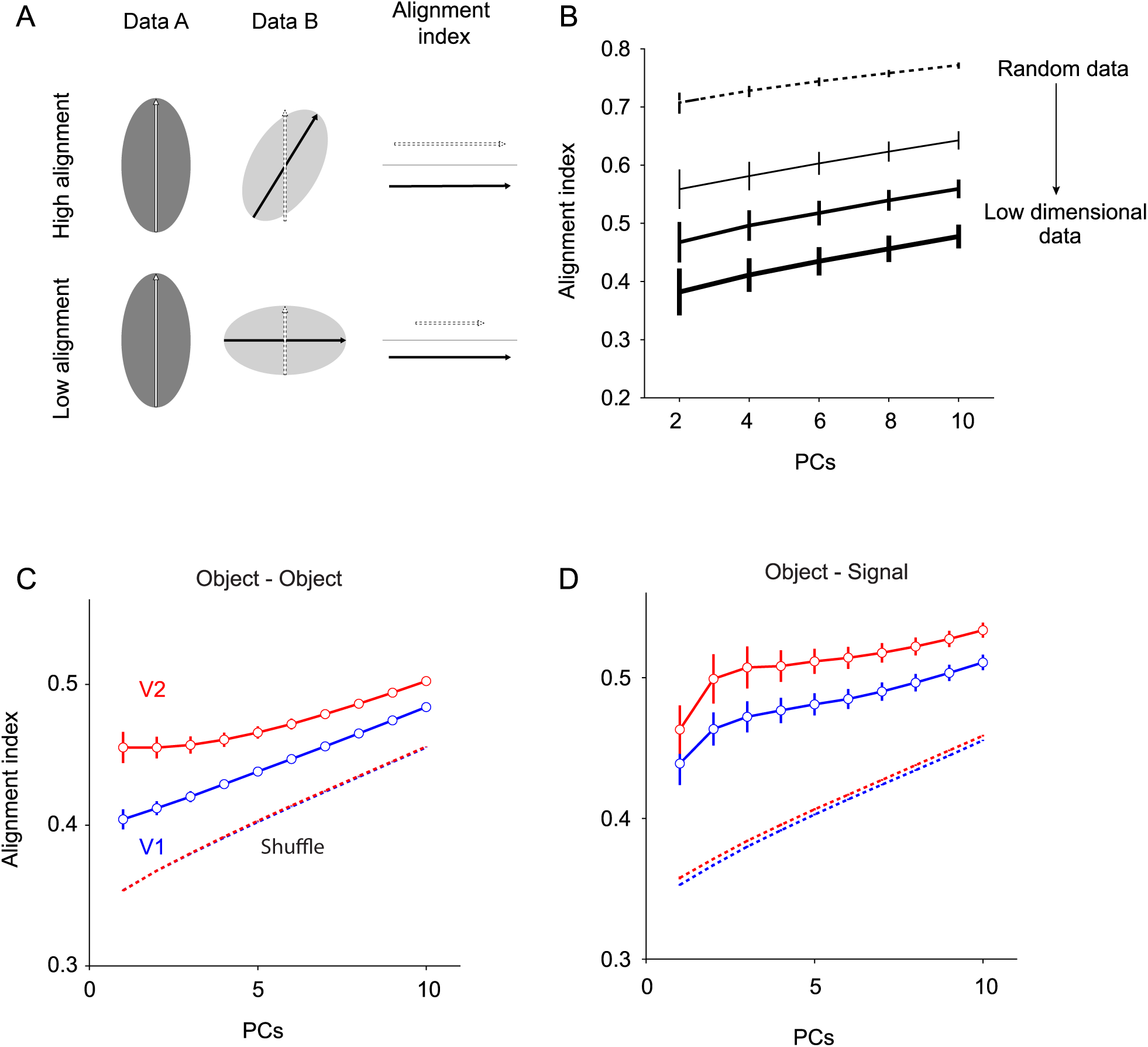
Subspace alignment. (A) Schematic showing how the alignment index is computed. The principal components of the two data sets, A and B, are computed (white and black arrows in left and center columns, respectively). The variance of data B that is captured by the principal component of A (dotted arrow, middle column; length an indication of the projection of data B onto that dimension) is compared to the variance captured by the principal component of data B (black line). The alignment index is the ratio of these two quantities (right column). The alignment index can be computed for an arbitrary number of principal components. An example of high alignment is shown at the top; an example of low alignment at the bottom. (B) The alignment index for two sets of synthetic data, that vary from being fully random (dotted line) to low dimensional (thick line). (C) The alignment between the different object manifolds is higher in V2 (red) compared to V1 (blue). Shuffling the responses of the object manifold results in lower alignment in both areas (corresponding dotted lines). Error bars are SEM across the 45 object-object comparisons. (D) The alignment between the object manifold and the signal manifold is higher in V2 compared to V1. The alignment decreases when we shuffled the responses of the object manifold. The error bars are the SEM across the 10 object-signal comparisons.

To illustrate the behavior of this metric, we first applied it to synthetic data: responses sampled from random, Gaussian distributions. The alignment score between two such random distributions is high (Figure 4B, dotted line). Note that for high-dimensional responses this occurs not because the principal components for the two data sets are similar, but because nearly all principal components explain the data equally well. Thus, the alignment score will be 1 even if the identified PCs are quite different from each other. We then generated low dimensional response distributions, by scaling the variance along the first few dimensions (see Methods). As the data became lower dimensional (Figure 4B, thicker line), the alignment score dropped. This is because with more prominent structure in the data (i.e. variance higher along some axes than others), it is unlikely that the leading principal components for one data set will capture substantial variance of another random data set, unless the structure of those responses is similar.

We computed an alignment index for population responses to each pairing of the 10 object manifold responses (45 object-object manifold comparisons). The alignment index between object manifolds was higher in V2 than in V1 (Figure 4C, red compared to blue line; p<0.001 for each comparison, paired t-test). Importantly, the alignment index is higher in V2 although the response distributions there were lower dimensional than in V1 (Figure 2). For comparison, we also calculated the alignment after shuffling responses across samples, which eliminates the low-dimensional structure in the responses. In this case, the alignment indices were similar in V1 and V2 (dashed lines, Figure 4C), and significantly lower than the indices for the measured responses. This indicates that different object manifolds have shared structure in both V1 and V2, but the degree to which those dimensions are similar across textures is higher in V2.

Higher alignment across the many different pairings of object manifolds in V2 would suggest a global organizational structure across textures which is more prominent in V2 than V1. We tested this possibility by measuring the alignment of each object manifold with the signal manifold (i.e. the alignment of each of the 10 object manifolds with the signal manifold). The alignment index was also higher in V2 than V1 (Figure 4D; p<0.05 for comparisons of 3 PCs or more; Wilcoxon signed-rank test). Shuffling the object manifolds decreased their alignment with the signal manifold in both V1 and V2 (dotted lines, Figure 4D).

We conclude that the representation of textures of V2 is a more structured response than in V1, in which the responses to different textures and samples of those textures are aligned in a low-dimensional embedding.

### Implications for discriminability and generalizability

The differences between the representational geometries in V1 and V2 might be expected to influence discriminability between different textures. We first tested the consequence of the alignment of the object manifolds to each other (and to the signal manifold), which was higher in V2 than V1. The greater alignment indicates that the variance in population responses attributable to different samples lies at least partly in the axes defined by responses to different textures. This alignment might thus be expected to reduce the discriminability between different textures. To test this possibility, we shuffled the responses across different samples for each neuron separately. Shuffling destroys the object manifold structure, resulting in spherical response distributions. If the primary axes of variance of the object manifold are aligned with each other, as our analysis indicates, then discriminability should improve after shuffling (Figure 5A, top). In contrast, if the primary representation axes are misaligned, shuffling would reduce performance (Figure 5A, bottom).

**Figure 5:**
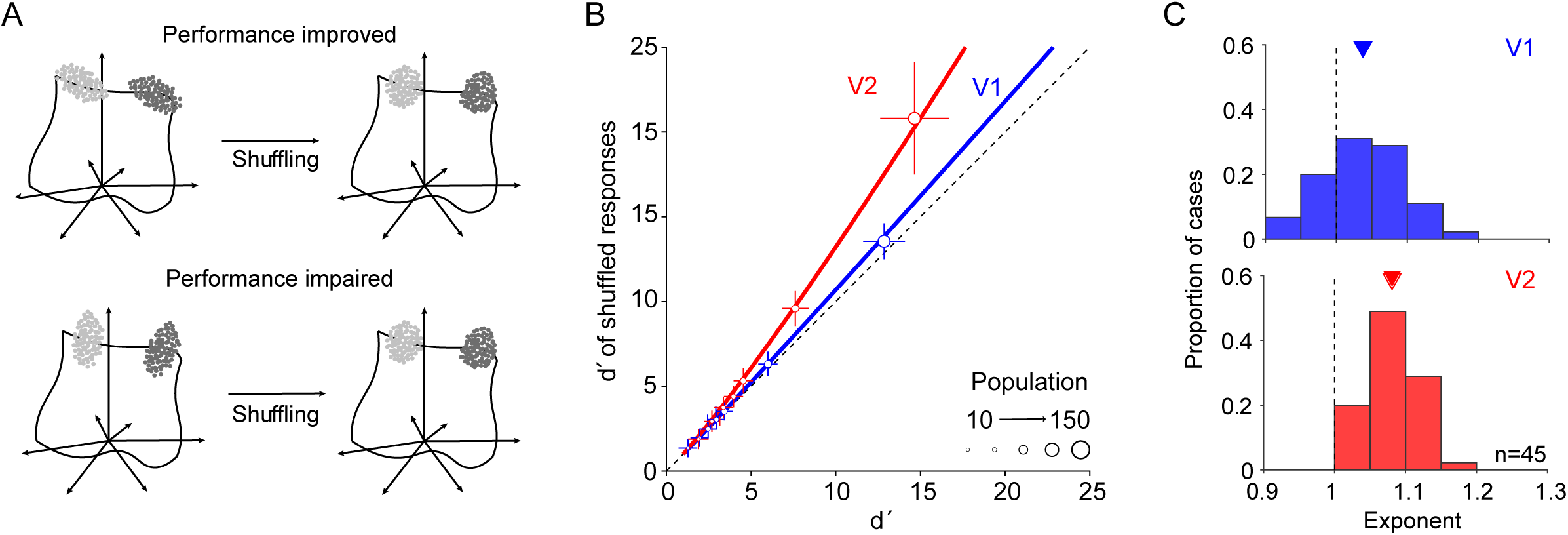
Testing the alignment of the object manifolds. (**A**) Top: If the variances of the object manifolds (light and dark gray clouds) are aligned with each other, then shuffling (right) will improve performance. Bottom: In contrast, if variance is highest in a direction misaligned with the axis between the centroids of the two object manifolds, then shuffling should increase the overlap between the response distributions. (**B**) A comparison of the d’ values after (ordinate) and before (abscissa) shuffling, for V1 (blue) and V2 (red) populations of different sizes (symbol size) for one of the 45 pairwise comparisons of object manifolds. Error bars are the SD over the 200 different subsamples for each population size. The solid line is the power law fit. (**C**) Distribution of power law exponents for V1 (top) and V2 (bottom) for each of the 45 pairwise discriminations of object manifolds.

We assessed the effect of shuffling responses on the discriminability of each pairing of object manifolds (n=45 cases), for populations of different sizes. Figure 5B compares the performance after shuffling (ordinate) with the performance for the original data (abscissa) for an example pairing of object manifolds, for populations of differing size (symbol size). Shuffling resulted in better discriminability in both V1 (blue) and V2 (red), but the effect was stronger in V2.

To quantify this difference, we fit a power law function separately to the performance curves (d’ of the shuffled compared to unshuffled data) of each of the 45 object manifold discriminations. This function provided excellent fits to the data: the mean variance accounted for was 97.7% and 99.5% in V1 and V2, respectively. The mean exponent of the function was higher in V2 (Figure 5C, red; 1.081±0.006 SEM; p<0.001, t-test for difference from 1) than V1 (1.038±0.009, p<0.001, t-test; for the comparison of the two areas, p=0.004, paired t-test), indicating a stronger improvement in performance with shuffling.

A second implication of building a systematic, low-dimensional representation of textures is that this low-dimensional space will be useful for discriminating between each pair of textures, even those that were not used to estimate the dimensions of the space.

To test this possibility, we compared discriminability afforded by responses to two object manifolds in the subspace defined by the signal manifold. We first performed PCA on the responses of 50 randomly chosen V1 or V2 neurons to the signal manifold stimulus ensemble. We then projected responses of two object manifolds into the subspace spanned by varying numbers of the identified principal components. Importantly, the responses to the object manifold stimulus ensemble (and the texture families which elicited them) were not used to define the signal manifold subspace in which the decoding was done.

The discriminability between two example object manifolds, in V1 (Figure 6A, blue solid line) and V2 (Figure 6B, red solid line), improves as the dimensionality of the subspaces increases, as expected. In V2, performance in the subspace defined by the first few principal components of the signal manifold is better than in a space spanned by random vectors, defined by performing the same analysis after shuffling the signal manifold responses in each population (black line). In V1, the difference between the signal manifold subspace and the shuffled space is smaller, and performance in the first few principal components of the signal manifold is lower than in V2.

**Figure 6:**
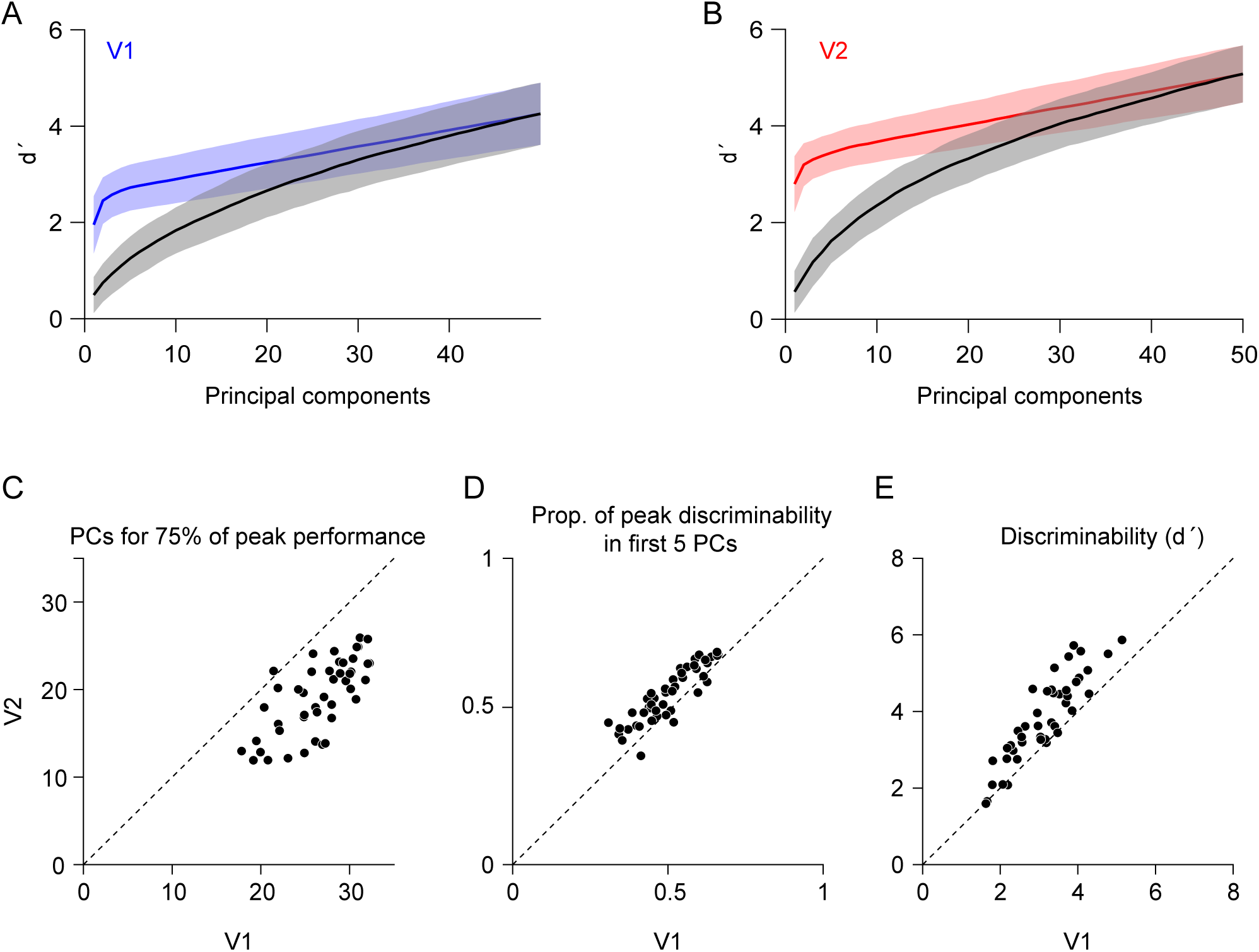
Signal manifold dimensions allow for better discriminability between novel textures from the object manifold stimulus ensemble. (**A**) The d ՛ values for V1 responses to two textures of the object manifold ensemble (ordinate), after projecting onto signal manifold subspaces of different dimensionalities (abscissa). Dotted line represents projections onto the subspaces after shuffling (across stimuli) responses to the signal manifold ensemble. (**B**) Corresponding data from V2. (**C**) A comparison of the number of PCs needed to achieve 75% of the performance of the full population, in V1 and V2. Fewer dimensions are needed in V2. Points are slightly jittered to reduce overlap, resulting in non-integer values of dimensions. (**D**) Proportion of the discriminability (d ՛) of the full population that is achieved by projecting the object manifolds onto the first five dimensions of the signal manifold, in V1 and V2. A higher proportion is achieved in V2. (**E**) Discriminability (d ՛) between object manifolds in the full population space. V2 responses afford better discrimination.

To aggregate performance across the 45 different pairings of object manifolds, we extracted several measures from these performance curves. First, we measured the number of dimensions needed to capture at least 75% of the discriminability evident in the full 50-neuron population space (Figure 6C). The number of dimensions needed was consistently higher in V1 than V2 (26.6±0.6 SEM vs 19.6±0.6; p<0.001, paired t-test). Second, we measured how much of the peak discriminability was achieved by projecting responses onto the first 5 principal components of the signal manifold. The proportion was consistently higher in V2 than V1 (Figure 6D, 0.55±0.01 vs 0.50±0.02, p=0.008, paired t-test). Finally, we compared the discriminability afforded by the V1 and V2 populations: discriminability was consistently better in V2 (Figure 6E, mean d ՛ of 3.82±0.16 vs 3.19±0.013, p<0.001, paired t-test). Notably, although peak performance was higher in V2, near peak (75%) discriminability was reached in a lower dimensional space (Figure 6C) and a higher proportion of peak discriminability was evident along the first few principal components (Figure 6D).

### Representational geometry in convolutional neural networks

Our results show a change in the representational geometry for textures between V1 and V2. We wondered whether a similar evolution would be evident in hierarchical, convolutional neural networks (CNNs), whose organizations have some similarities with the primate visual system (Yamins and DiCarlo, 2016; Lindsay, 2021). For instance, the responses of individual neurons in the visual cortex can be well-modeled as linear combinations of units in CNNs trained on object recognition tasks (e.g., Cadena et al., 2019; Bashivan et al., 2019). However, whether the representational geometry of these artificial networks evolves across layers in a similar manner to the changes we observe between the first two stages of cortical visual processing is unknown.

We analyzed the representational geometry of two widely-studied hierarchical CNNs—AlexNet and VGG16. We first measured how the response dimensionality for textures changed across layers of the network. We considered only responses in the convolutional, rectification (‘ReLU’), and pooling layers, ignoring the layers including and after the first fully connected layer, because the architecture and representation in these layers is distant from early visual cortex. We randomly selected 50 units in each layer of each network and analyzed the responses of these units to the same signal and object manifold ensembles we had used in our cortical recordings.

In the visual cortex, the participation ratio for neuronal population responses to textures was lower in V2 than V1 (Figure 2). In AlexNet, the participation ratio for responses to the signal manifold ensemble also dropped strongly after the first convolutional layer (Figure 7A, black), but then showed either a maintained or slightly increasing dimensionality. There was no consistent change in the ReLU or pooling layers (Figure 7A, brown and tan). In VGG16, the convolutional layers showed a decrease in participation ratio, particularly in the first few layers (Figure 7B, black; Spearman correlation, r=-0.57, p=0.05), but there was a consistent increase in the participation ratio across pooling (r=1, p=0.02) and ReLU layers (r=0.93, p<0.001). For the object manifold, there was no consistent reduction in the participation ratio across layers in either AlexNet (Figure 7C) or VGG16 (Figure 7D), expect for a slight but consistent increase across ReLU layers of the former (r=0.68, p=0.01).

**Figure 7:**
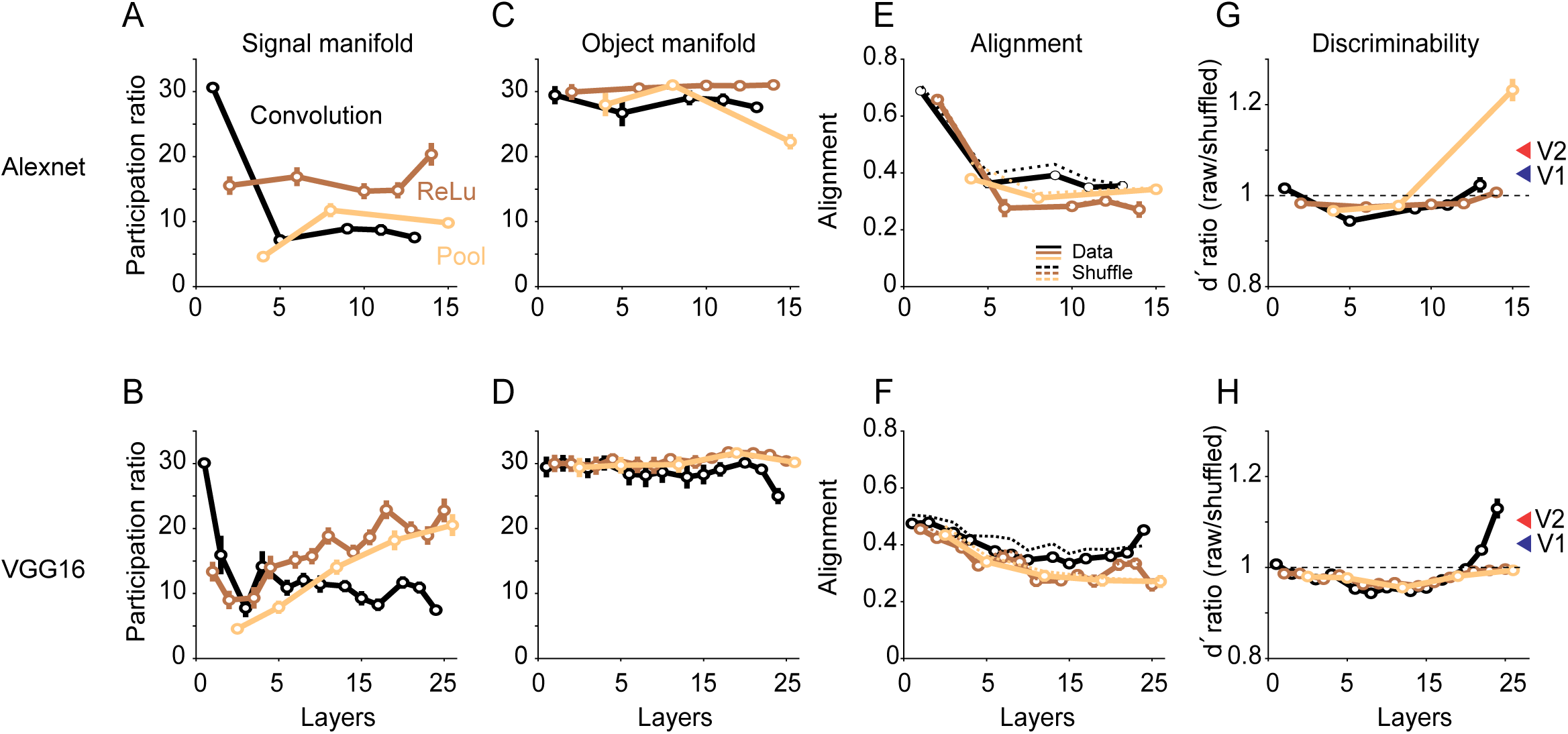
The representational geometry of Convolutional Neural Networks (CNNs) for textures. (A) The participation ratio for responses of 50 units in AlexNet to the signal manifold texture ensemble, in different network layers. Layers are grouped by their type: convolutional layers (black), ReLU layers (brown), and pooling layers (tan). Error bars indicate standard deviations across 200 different samples of units. (B) The analysis of (A) but for VGG16. (C) Participation ratio for the responses to the object manifold texture ensemble, following the conventions of (A). Error bars represent the SEM across the 10 object manifold ensembles. (D) The analysis of (C) but for VGG16. (E) The alignment between each pairing of object manifold responses, for each layer of AlexNet. Alignment between the object manifolds after shuffling the responses across texture samples are shown in the thin lines. (F) The analysis of (E) for VGG16. (G) Comparison of discriminability between responses to two object manifold ensembles, afforded by raw responses of 50 AlexNet units compared to the shuffled responses. Shuffling generally reduces performance (ratios less than 1), unlike the effect for V1 and V2 responses, shown by the blue and red arrowheads respectively. Arrowheads represent the mean effect observed in V1 and V2 for populations of 50 neurons (ratio of the quantities plotted in Figure 5, for populations of 50 neurons). Error bars represent the SEM across the 45 different pairings of the object manifold responses. (H) Analysis of (G) for VGG16.

In cortex, the alignment of different object manifolds to each other was higher in V2 than V1 (Figure 4). In contrast, the alignment of the object manifolds in AlexNet decreased systematically across the first layers before reaching an asymptote (Figure 7E); in VGG16, there was also a consistent decrease in alignment across both ReLU (Figure 7F, brown; r=-0.73, p=0.007) and pooling (tan; r=-1, p=0.02) layers. Notably, the alignment of responses in both networks was lower than alignment of shuffled responses (Figure 7E,F, dotted lines), whereas the cortical responses were more aligned than the shuffle controls.

Finally, we tested how shuffling responses within the object manifolds affected discriminability. In the neuronal responses, shuffling increased discriminability, particularly in V2 (Figure 5, replotted in Figure 7G,H in arrowheads, as the ratio of d ՛ values for shuffled vs raw data). In contrast, in both networks the discriminability for shuffled responses was generally similar to or worse than the discriminability for raw responses (ratios less than 1, indicated by dotted line; Figure 7G,H), except in the final few layers of the network. The discriminability ratios were significantly smaller (p<0.001, bootstrap test) than the ratios for the neural data in both networks, except in the final layers of each network. We conclude that the representational geometry of these networks, as assessed by our metrics, differs from that in cortex.

## Discussion

Using Neuropixel recordings in the macaque visual cortex, we compared the representational geometry of V1 and V2 neuronal population responses for textures. We draw three primary conclusions. First, in V2 there is an emergence of a lower-dimensional representation of texture images, relative to V1. This is evident in the lower participation ratios and the better alignment of different object manifolds to each other and to the signal manifold, in V2 compared to V1.

Second, the representation in V2 affords better discrimination between novel, out-of-set textures. Third, the changes in representational geometry we observe between V1 and V2 are not evident in common CNNs, similar to those that have been used successfully to model single neuron selectivity in early cortex. Thus, our results suggest that comparisons between the response geometry of networks and neuronal populations can reveal differences in representational strategy not evident from assessing single neuron tuning.

### Representational geometry

Several recent studies have studied visual representations from a geometric perspective, as we have done, including work in macaque V1 (Cowley et al., 2016) and IT (Lehky et al., 2014), and in mouse V1 (Stringer et al., 2019; Froudarakis et al., 2021) and somatosensory cortex (Nogueria et al., 2023).

Particularly relevant to our study, Stringer et al. (2019) measured the dimensionality of neuronal population representations in mouse V1 to 10,000 natural images. They characterized the representational geometry by fitting a power law to the eigenspectrum of the measured responses. The variance captured by the nth eigenvector scaled as 1/n, which the authors argue indicates a representation which balances the need for a smooth, differentiable representation with a high-dimensional, efficient code. For comparison, we applied the analysis of Stringer et al. (2019) to our data and found that the eigenspectra for V2 responses to textures were fit with a larger exponent (indicating a lower dimensional code) than the V1 responses (Supplementary Figure 2), consistent with the conclusions reached by our measurements of the participation ratio.

Our study extends this prior work in two important ways. First, we provide a comparison of representational geometry across stages of processing, rather than attempting to infer the coding strategy from the analysis of population responses in a single area and comparison to an external model or theoretical ideal. A comparative approach has previously been used to explore how single neuron response statistics evolve across stages of cortical processing, including measurements of lifetime sparsity (Willmore et al., 2011) and neuronal tolerance and selectivity (Rust and DiCarlo, 2010), among others. Our approach reveals that the representation of textures but not gratings becomes lower dimensional between V1 and V2.

Second, we provide an assessment of the representational geometry that extends beyond a simple quantification of dimensionality. We compare the alignment of different axes of the representation with each other (Figure 4) and relate our inferences of the geometry to stimulus discriminability (Figure 5 and 6). We posit that such efforts are critically needed. Comparisons of dimensionality are a first step but provide limited understanding of the nature of the representation. Characterizing the geometry and its functional implications will require numerous complementary metrics.

Our focus on texture stimuli to study representational geometry in V1 and V2 was motivated by prior work showing that V2 neurons differentiate between synthetic naturalistic textures and spectrally-matched noise images, whereas V1 neurons do not (Freeman et al., 2013; Ziemba et al., 2016; see also Bolanos et al., 2024 for related work in mouse visual cortex). The emergence of sensitivity to texture statistics in V2 indicates a notable transformation in the representation of these images between V1 and V2. Our observations of differences in the representational geometry for these stimuli in V1 and V2 is consistent with this view, though obviously different in its focus on population-level representation across textures.

Neither prior single-neuron studies nor our study reveals which specific statistics are encoded in V2. Because texture statistics are high dimensional (458 parameters for the textures used here), inferring single neuron selectivity can be cumbersome. Our observations suggest a worthwhile alternative approach might to relate the primary representational axes in V2—the latent dimensions—to the parameters of the texture synthesis algorithm, to infer the stimulus properties that define the representation (Chang and Tsao, 2017). In this view, understanding the transformations between stages of processing could be conceptualized as the computations needed to change the representational geometry in a particular way (Kriegeskorte and Kievit, 2013), as opposed to the computations needed to generate a particular receptive field property evident in individual neurons.

If V2 neurons encode higher order texture statistics, why do we find that object manifolds are slightly lower dimensional in V2 than V1? In terms of Portilla and Simoncelli (2000) statistics, the different samples that make up the object manifold are nominally identical. We see two possibilities. First, though the images are synthesized to have matched statistics (on average, across space), the image components that fall within the receptive field (and surround) of each neuron are not identical (Ziemba and Simoncelli, 2021). Thus, the same encoding that generates the structure of the signal manifold will lead to similar (though weaker, as observed) structure in the object manifolds. A second possibility is that the different samples differ in statistics other than those considered by the Portilla and Simoncelli algorithm and that V2 neurons are more sensitive to these statistics than V1 neurons. In any case, our data clearly show that different samples of synthesized textures modulate neuronal responses, particularly in V2, in a manner consistent with (but weaker than) the modulation induced by different textures.

### Caveats and limitations

Our recordings were performed in sufentanil-anesthetized monkeys, which might have affected the measured responses. However, we note that differences in texture selectivity were first documented in this preparation (Freeman et al., 2011; Ziemba et al., 2016) and later validated in awake animals, although there are some differences in the dynamics of selectivity (Ziemba et al., 2024). Thus, we expect the geometries that we measure would be similar in awake animals, though this will need be tested experimentally.

Our estimates of representational geometry were based on limited numbers of neurons and images, reflecting standard experimental challenges of presenting many images to many neurons. The sampling required to accurately measure geometry in high dimensional data (i.e. populations of many neurons) is an area of ongoing research, but undersampling can lead to systematic biases in the estimation of dominant structure (e.g., Altan et al., 2021; Landau et al., 2023; Pospisil and Pillow, 2024). Our results could be affected by these confounds. On the other hand, the comparisons we make are between identically sized populations in V1 and V2, recorded in the same animals. Thus, we expect that the differences we observe between V1 and V2 should be robust to these issues, even if some quantities are misestimated because of limited sampling.

We analyzed the measured neuronal population responses using simple, linear dimensionality methods, consistent with previous studies of visual cortex (Lehky et al., 2014; Cowley et al., 2016; Stringer et al., 2019). Application of non-linear methods could reveal additional differences (or similarities) between the representations of V1 and V2. Non-linear dimensionality reduction methods have revealed important topological structure in mouse hippocampus (Nieh et al., 2021) and the anterodorsal thalamic nucleus and subiculum (Chaudhuri et al., 2021).

### Comparison to artificial networks

CNNs have emerged as a helpful tool for understanding cortical visual processing (Yamins and DiCarlo, 2016). There are notable similarities between the response properties of single neurons in V4 and IT and units in deep layers of these networks (Yamins et al., 2014). Linear combinations of filter responses can predict neuronal responses to diverse image ensembles in V1 (Cadena et al., 2019), V4 (Bashivan et al., 2019), and IT (Yamins et al., 2014; Ponce et al., 2019; Bao et al., 2020). These observations suggest similarities in the representations of artificial and biological networks.

In contrast, we found notable differences in the representational geometry of visual cortex and those in CNNs. Whereas there was a reduction in response dimensionality and the emergence of more shared structure in V2 compared to V1, this evolution was not prominent across layers of AlexNet or VGG16. The alignment of the different object manifolds with each other was entirely absent in the CNNs. Thus, though CNNs may replicate some aspects of single neuron tuning, the population representation deviates strongly from that seen in the brain, at least for the metrics that we consider.

Similar discrepancies have been noted by some but not all population-based comparisons of visual cortical and CNN representations. For instance, representational similarity analysis (RSA; Kriegeskorte et al., 2021) applied to human fMRI responses to natural and artificial images revealed clear disparities with the representations of 14 different CNNs in some (Xu and Vaziri-Pashkam, 2021) but not other studies (Khaligh-Razavi and Kriegeskorte, 2014). For neuronal population responses, RSA suggests similarities between representations in IT and those seen in deep layers of CNNs (Yamins et al., 2014). Though RSA provides an attractive framework for comparing representations, particularly those measured with different experimental approaches, it provides limited functionality for understanding the basis of a particular similarity score (Dujmovic et al., 2024). An alternative approach to characterizing population representations is to calculate the representational ‘capacity’, a metric which, as the name suggests, emphasizes properties that support classification or discriminability performance. The representational capacity depends on manifold dimensionality and radius and on correlations across manifolds. These metrics evolve similarly across stages of networks and hierarchically organized cortical areas of the mouse (Cohen et al., 2020; Froudarakis et al., 2021; Bolanos et al., 2024).

The difference in the aspects of representational geometry we consider between the visual cortex and CNNs could arise from any of the many obvious differences between the two—their architecture, training diets, and so on. But recent work suggests a key factor may be the training objective, which can strongly influence the representational geometry. The CNNs we analyzed were trained to support objection classification. The visual system must of course support object classification (or recognition) as well, but it must also allow us to generalize knowledge across instances and contexts and to be robust to noise. These desiderata may be better accommodated by low dimensional representations: when artificial networks are trained on multiple tasks, or when confronted with noise or adversarial images in training, they tend to build lower dimensional representations (Johnston and Fusi, 2023). Notably, though the representation in V2 for textures is lower dimensional than in V1, there is no loss of discriminability between textures: neuronal populations in V2 afforded better absolute performance for discrimination between pairs of textures. It thus appears that the cortical representation might build more generalizable, noise robust representations across stages of processing while maintaining discriminability between stimuli.

## Methods

### Experimental model and subject details

Animal procedures have been described in detail in previous work (Smith and Kohn, 2008). Briefly, animals (macaca fascicularis, male 2-4 years old) were anesthetized with ketamine (10 mg/kg) and maintained on isoflurane (1%-2%) during surgery. Recordings were performed under sufentanil (typically 6-24 μg/kg/hour) anesthesia. Vecuronium bromide (150 μg/kg/hr) was used to prevent eye movements. Refraction was provided by supplementary lenses. Data from one animal were excluded from the study, though its inclusion would not alter any of the conclusions provided, because proper refraction was elusive. The duration of the experiment varied from 5-7 days. All procedures were approved by the IACUC of the Albert Einstein College of Medicine.

### Visual stimulation and recordings

Recordings were performed using the Neuropixel 3B probes (IMEC). Probes tips were sharpened using a grinding wheel and then mounted on custom holders. Probes were lowered into the cortex using a microdrive, attached to a Kopf micromanipulator. As we performed a full durotomy over the opercular cortex, no guide tube was used during insertion. Data was acquired on the first 384 interleaved channels of each probe (spanning ∼7.6 cm), using SpikeGLX software at a sampling rate of 30 KHz. The amplitude gain was adjusted for each probe to prevent clipping. A high pass filter with a cutoff frequency of 300 Hz was used. Kilosort 2.5 was used to perform an initial, automatic spike sorting, followed by manual curation using Phy. Stimuli were presented on a calibrated CRT monitor placed 70-95 cm from the animal (1024 x 768 pixel resolution, 100 Hz frame rate, ∼ 40 cd m^-2^ mean luminance).

In each recording session, we first measured the spatial receptive fields by briefly presenting small (1.2 deg diameter) drifting gratings, at a range of spatial positions. We used the resultant receptive field maps to center our stimuli on the aggregate receptive fields of the recorded neurons and to discern the transitions between cortical areas.

We obtained textures from online databases (Brodatz texture database, Abdelmounaime and Dong-Chen, 2013; Kylberg database, Kylberg, 2011; Oulu texture database, Lee and Bang, 2019; and the Salzburg texture images, https://www.wavelab.at/sources/STex/). We used the Portilla-Simoncelli algorithm (Portilla and Simoncelli, 2000) to generate different synthetic textures whose statistics matched those of each chosen source texture. Each texture was matched for mean luminance (128 on an 8 bit, 0-255 range) and RMS contrast (0.5). The textures were 256 x 256 pixels in size, corresponding roughly to 6 x 6 degrees (40-44 pixels/deg, depending on viewing distance). All textures were presented in a pseudo-random order. Each stimulus was presented for 200 ms with an inter-stimulus interval of 200 ms.

In a subset of recordings, we presented ensembles of drifting gratings. Gratings (3 or 4 cyc/s, full contrast) were presented in a circular aperture 8-12 degrees in diameter for 200 ms with an inter-stimulus interval of 200 ms. We presented 324 distinct gratings, consisting of all combinations of 18 different orientations (0 to 170 degrees, in increments of 10 degrees) and 18 spatial frequencies (0.04 to 16 cyc/deg), in increment of 0.5 octaves. Each grating was presented 50 times. To compare the dimensionality of population responses evoked by these stimuli, we randomly selected different random subsets of 90 gratings, to match the number of stimuli used in the texture experiments

### Data Analysis

We measured firing rate in a 200 ms window beginning 40 ms after stimulus onset, to account for response onset latency. We subtracted the spontaneous firing rate, measured in a 50 ms window beginning 30 ms before stimulus onset and extending 20 ms after, from the measured firing rate. To consider only neurons that were visually responsive to the presented stimuli, we discarded any neurons that fired less than 1 spike/second on average (across stimuli), for a given stimulus ensemble. We also discarded neurons for which the average response across stimuli was not statistically distinguishable from the spontaneous response (p≤0.05, ANOVA). Finally, we compute the geometric mean of the Fano factor (of the spike counts) across all stimulus conditions for each neuron and discarded any unit with a value above 3.5. This final selection criterion was done to remove units whose responses were unreliable, which in many cases we attributed to a failure to maintain spike isolation for the duration of the recording.

We performed principal component analysis on the measured V1 and V2 responses. For measurements of the signal manifold, we averaged responses across the 10 presentations of each unique image and the 10 samples of each texture and performed PCA on the responses to the 90 distinct textures. For measurements of the object manifold, we averaged across the 10 trials and performed PCA separately on the responses to each of the 10 object manifold ensembles (each with 90 samples). In both analyses, we z-scored the responses of each neuron across images before performing PCA. We analyzed population responses of 10, 20, 30, 40, 50, 100 or 150 neurons, chosen randomly without replacement from the set of sampled units. We analyzed 200 separate random subpopulations.

We used the participation ratio to quantify the dimensionality of each population response. If *λ_i_* are the eigenvalues of the principal component decomposition of the response covariance, then the participation ratio, *PR*, is defined as:

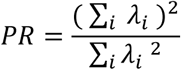

The alignment between a reference data set *A* and a test data set *B* was defined using the alignment index of Elsayed et al., 2016. The alignment index is a ratio:

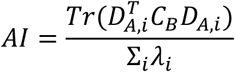

The denominator is defined by the sum of the top *i* eigenvalues of B. The numerator is defined as the variance of B that is captured by the top *i* PCs of data set A, which form *D_A,I_*, and *C_B_* which is the covariance of data set B. Essentially, the metric compares the response variance captured by PCs of a reference data set (in this, data case A) relative to the maximal variance that could be captured by that number of PCs of the data itself (in this case, test data set B).

Note that the alignment index is not symmetric: the alignment of test data A to reference data B need not equal the alignment of B to A.

To compute the alignment between object manifolds, we defined the role of test and reference randomly for the two object manifolds. To the compute the alignment between each object manifold and the signal manifold, we defined the object manifold as the test data and the signal manifold as the reference. For each condition, we computed the alignment index for 200 different random subpopulations of 50 neurons. For reference, we also calculated the alignment index after shuffling responses for each neuron across images (samples for the object manifold and different textures for the signal manifold), followed by PCA.

To illustrate the behavior of the alignment index, we first applied it to synthetic data (Figure 4B). We constructed matrices of population responses (1000 trials of 100 neurons) by draws from a random Gaussian distribution. We calculated the alignment between two such population responses, as well as low dimensional version constructed by scaling the variance along the first PCs by 1.25-2-fold. In all cases, we z-scored the responses before performing PCA and calculating the alignment index.

We measured the discriminability between two stimuli, *a* and *b*, afforded by the measured population responses:

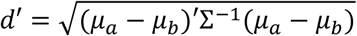

where 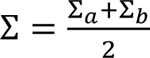, is the mean covariance of the population responses to *a* and *b*, with means of μ_a_ and μ_b_. We performed this analysis for populations varying from 10-150 neurons. We quantified the relationship between the d ՛ calculated from the measured responses and from shuffled data (shuffling across samples of the image manifold) by fitting a power law function, *d^’^_shuffled responses_=d^’^_measured responses_n*, where *n* is the reported exponent.

We used a similar approach for assessing the discriminability between different object manifolds, in the subspace defined by the signal manifold. We first performed PCA on the responses to the signal manifold ensemble. We then projected the population responses to two object manifold stimulus ensembles onto the subspace defined by increasing numbers of the identified principal components. We calculated d ՛ in these subspaces, using the approach described above.

We analyzed filter responses in pre-trained AlexNet and VGG-16 networks, using the same approach as applied to the neuronal data. We measured the participation ratio for responses of 50 units, chosen randomly, in each layer of each network, to the same stimuli used in the cortical recordings; we repeated this measurement for 200 different subsamples of units. We measured alignment between the first two principal components of the object manifold responses for each of the 45 pairings of object manifolds, using 200 different subsamples of 50 units. We measured the alignment between the object and signal manifold similarly. Finally, we measured d ՛ as for the neuronal data, using 200 different subjects of 50 units and measuring discriminability between each of 45 pairings of the object manifolds.

We used the bootstrap approach described by Efron and Tibshirani (1993) to assess the probability that the differences observed could have arisen by chance. Let *x*^→^ and *y*^→^ be vectors of samples (observations) with mean *x*_m_, *y*_m_ and standard deviation σ_x_ and σ_y_. We computed a t-statistic for the difference between these cases as:

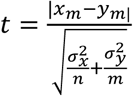

We then created two new vectors *x*^→^’ and *y*^→^’, where 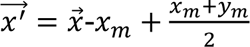 and 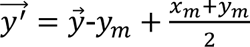.

We drew random samples without replacement, equal in number to the samples in the real data, from *x*^→^’ and *y*^→^’ and computed the same t statistic from these data. This process was repeated 1000 times and the p-value was defined as the location of the data in the rank ordered list of bootstrapped values.

## Acknowledgments

This work was supported by NIH awards EY030578, EY028626 and NS127107.

---

**Supplementary Figure 1:**
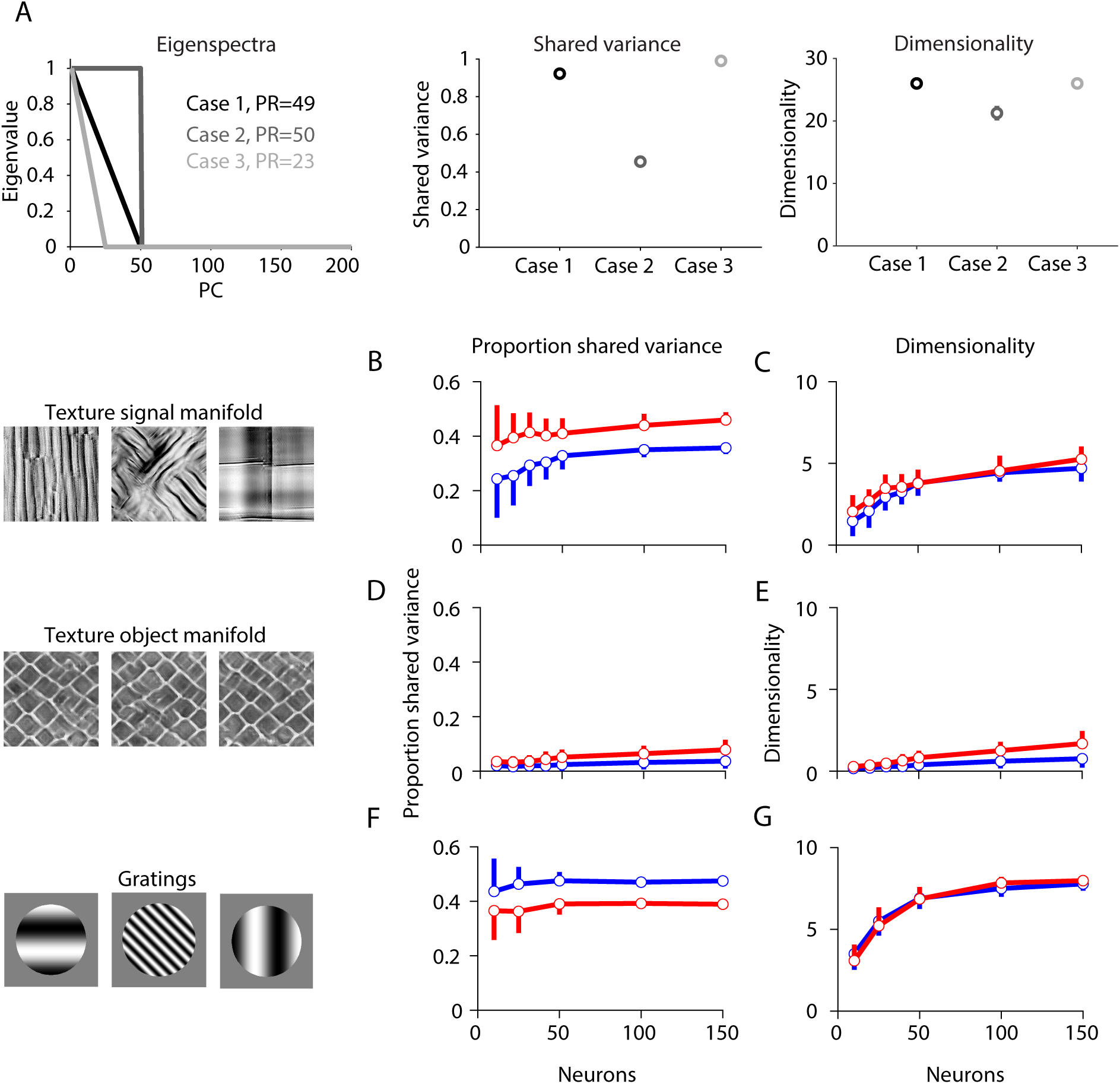
Representational geometry as measured by factor analysis. Factor analysis models the responses of a population of *n* neurons, *X*, as

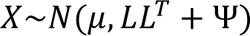

where μ is a vector of length *n* containing the mean responses of each cells, *L* is the loading matrix (of dimensions *n x m*) relating *m* latent variables to the neuronal responses and Ψ is a diagonal matrix. The matrix *LL*^T^ captures the shared covariance of the measured responses, whereas Ψ captures the variance that is private to each neuron. The parameters μ, *L*, and Ψ are estimated using expectation-maximization. We report two quantities that capture the properties of the factor analysis model fit to each data set (Williamson et al., 2016): the proportion of shared variance (defined as the ratio of *LL*^T^ to *LL*^T^ + Ψ) and the dimensionality (defined as the number of dimensions needed to explain 95% of the shared covariance, *LL*^T^). The latter is a more robust estimate of the dimensionality of the measured responses than the number of identified latent variables, *m* (Williamson et al., 2016; Semedo et al., 2019). (A) We conducted a simple simulation to illustrate how the factor analysis metrics we use—the proportion of response variance that is explained by the factors, and the dimensionality of the shared response covariance—complement the view provided by calculating the participation ratio for the PCA eigenspectrum. Left: The eigenspectra for 3 synthetic data sets, with the corresponding participation ratios. Note that Case 1 and 2 have, by construction, nearly identical participation ratios though the shapes of the eigenspectral differ. Middle: The shared variance identified by fitting a factor analysis model to these data. Note that Case 1 and 2 have strikingly different shared variance despite having similar participation rations. In addition, Case 3, which has a different participation ratio, has a similar amount of shared variance to Case 1. Thus, the participation ratio does not define whether the shared variance identified by factor analysis will be high or low. Right: The dimensionality of the shared response covariance, identified by factor analysis. Again, Case 1 and 2 have differing dimensionalities despite having similar participation ratios. Case 1 and 3 have similar dimensionalities despite having different participation ratios. (B) For the texture signal manifold stimulus ensemble, the shared variance identified by factor analysis is higher for V2 responses (red) than V1 responses (blue). The proportion shared variance in V2 is higher for each population size (p<0.001, bootstrap test). Error bars indicate standard deviation across 50 random subsets of neurons; only upper and lower error bars are shown due to overlap. (C) Dimensionality of the shared variance is similar but statistically distinct in V1 and V2 (p<0.05, except for population sizes of 50 and 100), and more shared dimensions can be identified in larger groupings of neurons. (D,E) Corresponding analysis for responses to the object manifold ensemble. Both the proportion of shared variance and the dimensionality are lower than for responses to different textures (signal manifold ensemble). V2 responses have slightly higher shared variance than V1 responses (p<0.05 for each comparison except populations of 20 and 30 neurons; bootstrap test). Error bars indicate SEM across the 10 different object manifolds. (F,G) Corresponding analysis for responses to the ensemble of gratings. Here, the proportion of shared variance is greater in V1 than in V2 (H; p<0.005 for all comparisons), opposite to the responses to textures. The number of shared dimensions is similar in the two areas (G).

**Supplementary Figure 2:**
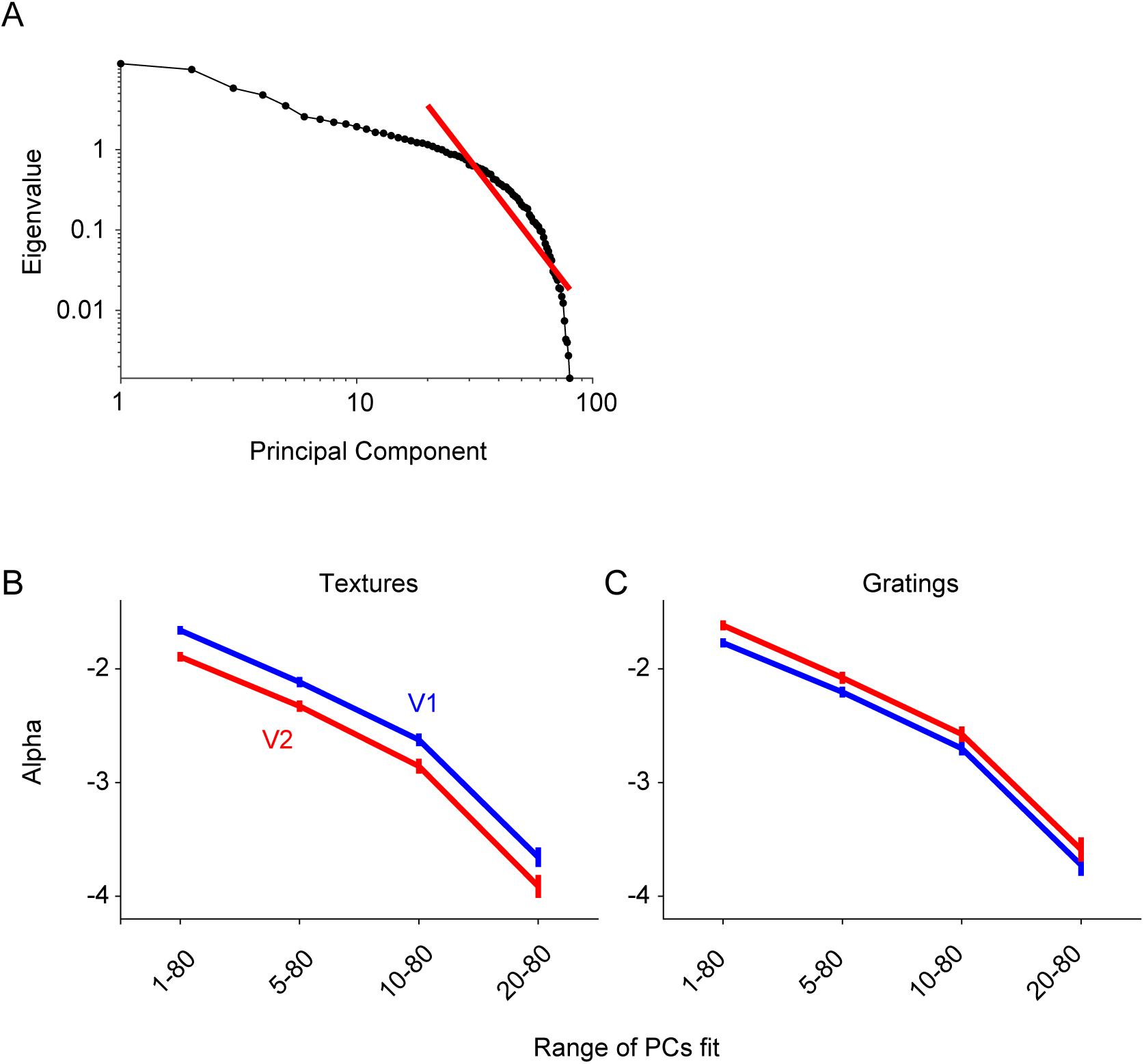
Assessing the representational geometry of V1 and V2 by measuring the decay of the eigenspectrum, as in Stringer, et al. (2019). (**A**) The eigenspectrum for the responses of population of 80 randomly selected V1 neurons to the grating stimulus ensemble (black), on log-log axes. The red line indicates a linear fit to the log-transformed data (i.e. a power law) for the 20^th^ to 80^th^ principal components. The slope of the line, or exponent of the power law, is −3.8. Note that the linear fit offers only a very rough approximation to the data. (**B**) The estimated slopes or exponents for the texture signal manifold ensemble, for V1 (blue) and V2 (red) populations of 80 neurons. We fit the spectra over different ranges of PCs, indicated on the abscissa. For each range, the slopes in V2 were significantly more negative than in V1 (p<0.001, bootstrap test), indicating a steeper drop off or lower-dimensional representation. To compute the fits, we randomly selected 80 neurons in each area (without replacement), performed PCA on the measured responses, and fit the resultant eigenspectra. We repeated the entire procedure 200 times. The indicated values correspond to the mean slopes; the error bars reflect the standard deviation of the distribution of 200 slope estimates. Larger ranges include more of the data, and in particular the first PCs which capture the most variance, but these data are less well approximated by the power law. We used a population size of 80 neurons in this analysis (as opposed to 50, as used in many other analyses) so that we could have a better assessment of the decay in the variance for higher PCs. (**C**) Corresponding analyses for responses to gratings. In this case, V1 slopes are significantly more negative than in V2 (p<0.001, bootstrap test). Though the power law fits to the decay of the eigenspectrum are tied to an elegant mathematical framework, the fits may be affected by estimation bias (Pospisil and Pillow, 2024). In addition, because our populations and image sets were smaller, our fits were to a smaller range of eigenvectors than the Stringer et al. (2019) spectra and the decay in this range was not well captured by a power law decay.

